# Bidirectional Modulation Of Synaptic Transmission By Insulin-Like Growth Factor I

**DOI:** 10.1101/2023.12.29.573627

**Authors:** José Antonio Noriega-Prieto, Laura Eva Maglio, José Carlos Dávila, Antonia Gutiérrez, Ignacio Torres Alemán, David Fernández de Sevilla

## Abstract

Insulin-like growth factor-I (IGF-I) plays a key role in the modulation of synaptic plasticity, and is an essential factor in learning and memory processes. Indeed, we have demonstrated that IGF-IR activation induces long-term potentiation (LTP) of synaptic transmission (LTP_IGF-I_) both in the barrel cortex, improving object recognition (Noriega-Prieto et al., 2021), and in the prefrontal cortex, facilitating the extinction of conditioned fear (Maglio et al., 2021). However, during aging, IGF-I levels are decreased, and the effect of this decrease in the induction of synaptic plasticity remains unknown. Here we show that the induction of NMDAR-dependent LTP at layer 2/3 PNs of the mouse barrel cortex is favored or prevented by IGF-I (10nM) or IGF-I (7nM), respectively, when IGF-I is applied 1 hour before the induction of Hebbian LTP. Analyzing the cellular basis of this bidirectional control of synaptic plasticity, we observed that while 10nM IGF-I generates LTP (LTP_IGF-I_) of the post-synaptic potentials (PSPs) by inducing LTD of the inhibitory post-synaptic currents (IPSCs), 7nM IGF-I generates LTD of the PSPs (LTD_IGF-I_) by inducing LTD of the excitatory post-synaptic currents (EPSCs). This bidirectional effect of IGF-I is supported by the observation of IGF-IR immunoreactivity at both excitatory and inhibitory synapses. Therefore, IGF-I controls the induction of Hebbian NMDAR-dependent plasticity depending on its concentration, revealing novel cellular mechanisms of IGF-I on synaptic plasticity and in the learning and memory machinery of the brain.

**SIGNIFICANCE STATEMENT:** Insulin-like growth factor-I (IGF-I) signalling plays key regulatory roles in multiple processes of brain physiology, such as learning and memory, and brain pathology, such as Alzheimer disease. Yet, the underlying mechanisms remain largely undefined. Here we demonstrate that IGF-I signalling triggers long-term potentiation (LTP) or long-term depression (LTD) of synaptic transmission at cortical synapses in a concentration dependent manner, thus regulating the induction of Hebbian synaptic plasticity. The present work represents an important conceptual advance in our knowledge of the cellular basis of IGF-I signalling in brain function.

## INTRODUCTION

Insulin-like growth factor-1 (IGF-I) is a peptide with well-known trophic functions in the brain, but its neuromodulatory role is still under study. It is well documented that IGF-I regulates neuronal firing (Carro et al., 2000; Nuñez et al., 2003; Gazit et al., 2016), and modulates synaptic transmission in several areas of the central nervous system (CNS), such as hippocampus or cerebellum (Nilsson et al., 1988; Araujo et al., 1989; Castro-Alamancos and Torres-Aleman, 1993; Seto et al., 2002; Fernando Maya-Vetencourt et al., 2012). IGF-I is also able to enhance glutamatergic synaptic transmission in rat hippocampal slices of juvenile animals through a mechanism that involves α-amino-3-hydroxy-5-methyl-4-isoxazolepropionic acid (AMPA) but not N-methyl-D-aspartate (NMDA) postsynaptic receptors, probably through phosphatidylinositol-3-kinase (PI3K) activation (Ramsey et al., 2005). However, in young adult and old rats, IGF-I significantly increases both AMPA and NMDA-mediated synaptic transmission by a postsynaptic mechanism (Molina et al., 2013). Additionally, the increase in expression levels of NMDA receptor subunits 2A and 2B at the hippocampus by IGF-I in aged rats (Sonntag et al., 2000) may facilitate the induction of long-term potentiation (LTP). Moreover, a reduction of inhibitory synaptic transmission dependent on the activation of the IGF-I receptor (IGF-IR) on astrocytes (Noriega-Prieto et al., 2021), and on the synthesis of nitric oxide (Noriega-prieto et al., 2022) has been demonstrated in the barrel cortex.

IGF-I is actively transported to the CNS from plasma through the blood-brain-barrier (Nishijima et al 2010), and is also locally produced by neurons and glial cells (Quesada et al., 2007; Rodriguez-Perez et al., 2016). Therefore, IGF-I levels in specific brain areas depend both on its local release and on its uptake from the circulation. Accordingly, the entry of serum IGF-I into specific brain areas is more significant during periods of increased neuronal activity or physical activity (Carro et al., 2000; Nishijima et al., 2010). Therefore, different brain areas may have different IGF-I levels depending on their levels of neuronal activity. Moreover, aging is associated with lower serum IGF-I levels (Breese et al., 1991) and impaired brain IGF-I activity (Muller et al., 2012), suggesting that IGF-I levels within as yet an undetermined range are essential for proper brain functioning. Thus, while brain IGF-I function varies according to age and brain activity, it is not known whether differing IGF-I concentrations will have distinct effects on the modulation of synaptic transmission, a key aspect of brain function.

In the present work, we have investigated the effect of different IGF-I levels in the modulation of the threshold of NMDAR-dependent LTP at layer 2/3 pyramidal neurons (L2/3 PNs) of the mouse barrel cortex. We demonstrate that 1 hour after application of 10nM IGF-I, Hebbian LTP is favored by reducing the number of pairings required to induce it, as well as the magnitude of the potentiation, whereas its induction is prevented after 7nM IGF-I. Analyzing the underlying effects, we found that 10nM IGF-I generates LTP of the PSPs (LTP_IGF-1_) by inducing LTD of inhibitory synaptic transmission and STP of the EPSCs. Conversely, 7nM IGF-I generates LTD of the PSPs (LTD_IGF-1)_ by inducing LTD of the EPSCs. Therefore, our results demonstrate that IGF-I levels are essential for the induction of Hebbian long-term synaptic plasticity at the barrel cortex by inducing a bidirectional long-term modulation of synaptic transmission, which may have important consequences in learning and memory processes in the mouse.

## MATERIALS AND METHODS

### Ethics statement and animals

All animal procedures were approved by the Ethical Committee of the Universidad Autónoma of Madrid, and Cajal Institute and are in accordance with Spanish (R.D. 1201/2005) and European Community Directives (86/609/EEC and 2003/65/EC), which promote the animal welfare. Male C57BL/6J mice were housed under a 12-h/12-h light/dark cycle with up to five animals per cage and were used for slice electrophysiology.

### Electrophysiological recording

Male C57BL6/J mice (12-18 days old) were decapitated, and brains were removed and submerged in cold (4°C) cutting solution containing (in mM): 189.0 sucrose, 10.0 glucose, 26.0 NaHCO_3,_ 3.0 KCl, 5.0 Mg_2_SO_4_, 0.1 CaCl_2_, 1.25 NaH_2_PO_4_.2H_2_O. Coronal slices (350 µm thick) were cut with a Vibratome (Leica VT 1200S), then slices were incubated (>1h, at room temperature, 25–27°C) in artificial cerebrospinal fluid (ACSF) which containing (in mM): 124.00 NaCl, 2.69 KCl, 1.25 KH_2_PO_4_, 2.00 Mg_2_SO_4_, 26.00 NaHCO_3_, 2.00 CaCl_2_, and 10.00 glucose). The pH was stabilized at 7.4 by bubbling the ACSF with carbogen (95 % O_2_, 5 % CO_2_). Slices were transferred to a 2 ml chamber fixed to an upright microscope stage (BX51WI; Olympus, Tokyo, Japan) equipped with infrared differential interference contrast video (DIC) microscopy and a 40X water-immersion objective and superfused at room temperature with carbogen-bubbled ACSF (2 ml/min). In some cases, picrotoxin (PiTX; 50 µM) was added to the ACSF to block GABA_A_-mediated inhibition. In these conditions, epileptiform activity was never observed in our sample.

Patch-clamp recordings from the soma of L2/3 PNs of the barrel cortex were performed in the whole-cell voltage-clamp and current-clamp configurations with patch pipettes (4–8 MC) filled with an internal solution that contained (in mM): 120 K-Gluconate, 10 KCl, 10 HEPES, 0.5 EGTA, 4 Na_2_-ATP, and 0.3 Na_3_-GTP, 10 NaCl buffered to pH 7.2–7.3 with KOH. (280 mOsm) Recordings were performed in the current- or voltage-clamp modes using a Cornerstone PC-ONE amplifier (DAGAN, Minneapolis, MN). Pipettes were placed with a mechanical micromanipulator (Narishige, Tokyo, Japan). The holding potential was adjusted to -60 mV, and the series resistance was compensated to ∼ 80 %. L2/3 PNs located over the barrels (layer 4) were accepted only when the seal resistance was >1 GΩ and the series resistance (10–20 MΩ) did not change (>10 %) during the experiment. Data were low-pass filtered at 3.0 kHz and sampled at 10.0 kHz, through a Digidata 1440A (Molecular Devices, Sunnyvale, CA). Synaptic responses were evoked with Pt/Ir concentric bipolar (OP 200 µm, IP 50 µm, FHC) stimulating electrodes (0.1 ms and 20-100 µA) connected to a Grass S88 stimulator and stimulus isolation unit (Quincy, USA). The stimulating electrode was placed at layer 4 of the barrel cortex. Single pulses (100-μs duration and 20-100 µA) were continuously delivered at 0.33 Hz. After recording 5 minutes of stable baseline of postsynaptic currents (PSCs), 5 minutes of postsynaptic potentials (PSPs) were recorded at 0.2 Hz. Then, the stimulation intensity was increased until ≈ 10% of the responses recorded during 5 minutes were suprathreshold. IGF-I was added to the bath and the recording was extended for 15 minutes. Then, we switched back to voltage-clamp, returning the intensity and frequency to that used for the baseline recording, and we recorded the PSCs during 20 minutes and the PSPs when PSC amplitude was stable. Then, we washed the IGF-I and we continuously switched between current-clamp and voltage-clamp to check the amplitude of the PSPs and PSCs after IGF-I washout.

The Intracellular solutions could also either contain: 1,2-Bis(2-aminophenoxy) ethane-N,N,N’,N’-tetra acetic acid (BAPTA; 20 mM). In some cases, IGF-I was not added and no changes either in synaptic transmission or in excitability were recorded (see supplementary Figure 1). The pClamp programs (Molecular Devices) were used to generate stimulus timing signals and transmembrane current pulses, and to record and analyze data. Plots of the changes induced by IGF-I on EPSC peak amplitude (percentage from control) versus time were constructed.

### Immunoelectron microscopy

After deep anesthesia with sodium pentobarbital (60 mg/kg), 6 month-old male mice (C57BL6/J) were perfused transcardially with 0.1 M phosphate-buffered saline (PBS), pH 7.4 followed by 4% paraformaldehyde, 75 mM lysine, 10 mM sodium metaperiodate in 0.1 M PB, pH 7.4. Brains were removed, post-fixed overnight in the same fixative solution at 4°C, coronally sectioned at 50 μm thicknesses on a vibratome (Leica VT1000S), and serially collected in wells containing cold PB and 0.02% sodium azide.

For IGF-IR immunogold labelling, sections containing the somatosensory cortex were used. Sections were first washed with PBS and incubated in a 50 mM glycine solution 5 minutes to increase antibody binding efficiency. Following a standard immunocytochemical protocol, tissue was first free-floating incubated in a rabbit polyclonal anti-IGF-IRα antibody (1/250; Santa Cruz) in a PBS 0.1M/1% BSA solution for 48 hours at 22°C. Then, sections were washed in PBS, and incubated with 1.4 nm gold-conjugated goat anti-rabbit IgG (1:100; Nanoprobes) overnight at 22°C. After post-fixing with 2% glutaraldehyde and washing with 50 mM sodium citrate, labelling was enhanced with the HQ Silver^TM^ Kit (Nanoprobes), and gold toned. Finally, immunolabeled sections were fixed in 1% osmium tetroxide, block stained with uranyl acetate, dehydrated in acetone, and flat embedded in Araldite 502 (EMS, USA). Selected areas were cut in ultrathin sections (70-80 nm) and examined and photographed with a JEOL JEM1400 electron microscope. As a control for the immunogold technique, sections were processed as above but omitting the primary antibody. No specific labelling was observed in these control sections.

### Data analysis

Pre- or post- synaptic origin of the synaptic plasticity induced by IGF-I was tested by analyzing the modification in the variance that parallel the synaptic current amplitude change, which reflect the change in transmitter release probability (Clements, 1990; Malinow et al., 1990; Kuhnt and Voronin, 1994) To estimate the modification of the synaptic current variance, we first calculated the noise-free coefficient of variation (CV_NF_) of the synaptic responses before (control conditions) and during IGF-I. We used the formula CV_NF_ = √ (δ_synaptic current_ ^2^ − δ ^2^)/*m*, where δ_noise_^2^ and δ ^2^ are the baseline and synaptic current peak variance, respectively, and *m* is the mean peak amplitude of the synaptic current. The ratio of the CV_NF_ (CV_R_) before (control conditions) and during IGF-I was obtained then for each neuron as CV_IGF-I_/CV_control_ (Clements, 1990) We constructed plots comparing variation in the normalized *m* (termed *M*) to the change in 1/CV ^2^) in each cell (Malinow et al., 1990) In these plots, depression of the synaptic currents has a presynaptic origin when values are below or in the diagonal, whereas points above the diagonal indicate a postsynaptic origin. However, potentiation of synaptic currents has a presynaptic origin when values are above or in the diagonal, whereas points below the diagonal indicate a postsynaptic origin (Faber and Kornt, 1991) This method requires a binomial EPSC amplitude distribution, a condition that must be met for the synaptic variance to reflect the probability of transmitter release. We could not directly test whether our data fitted a binomial distribution, but synaptic fluctuations were always evident and we assumed that synaptic release followed a binomial distribution. Data analysis was done in Clampfit 10 (Axon Instrument) and graphs were drawn in SigmaPlot 11. In all cases, statistical estimates were made with Student’s two-tailed *t*-tests for unpaired or paired data as required, and data are presented as means ± SE. The threshold for statistical significance was *P* < 0.05(*); *P* < 0.01 (**) and *P* < 0.001 (***) were also indicated.

## RESULTS

### IGF-I regulates the induction of Hebbian LTP (LTP_H_) in a concentration dependent manner

We have previously shown that 10 nM IGF-I decreases the induction threshold of LTP_H_ (Noriega-Prieto et al., 2021). Here, we analyzed the effect of a slightly lower dose of IGF-I on LTP_H_ induced by spike timing dependent plasticity (STDP) at layers 2/3 of the barrel cortex (**Fig. 1A**). We first checked in control conditions, the STDP protocol, consisting in a subthreshold PSP followed by a back-propagating action potential (BAP) at delays of 10 ms repeated 50 times at 0.2Hz (**Fig. 1B**) and then we performed similar experiments in slices in which previously IGF-I at doses of 10nM or 7nM was bath applied during synaptic stimulation of layer 4 at 0.2Hz (**Fig. 1C)**. In control conditions, the STDP protocol induced an LTP_H_ of PSPs (from 96.01 ± 4.12 to 137.62 ± 8.33 % of amplitude, P<0.001; n=7; **Fig. 1D and G, LTP control**). However, in slices pretreated with 10nM IGF-I, the STDP protocol induced a more robust LTP_H_ of PSPs (from 99.36 ± 1.08 to 141.59 ± 4.38 % of amplitude after 20 pairings, P<0.001; n=6 and from 99.36 ± 0.89 to 169.13 ± 6.36 % of amplitude after 50 pairings, P<0.001; n=6; **Fig. 1E and G, LTP facilitated**). Conversely, in slices pretreated with 7nM IGF-I, the STDP protocol did not induce LTP_H_ of the PSPs (from 101.66 ± 1.12 to 99.85 ± 6.35 % of amplitude after 50 pairings, P>0.05; n=6; **Fig. 1F and G, LTP impairment**). Thus, only 10 nM, but not 7 nM IGF-I favored the induction of LTP_H_. In other words, the activation of IGF-IRs favors or impairs LTP_H_ depending on IGF-I concentration.

**Figure 1.**
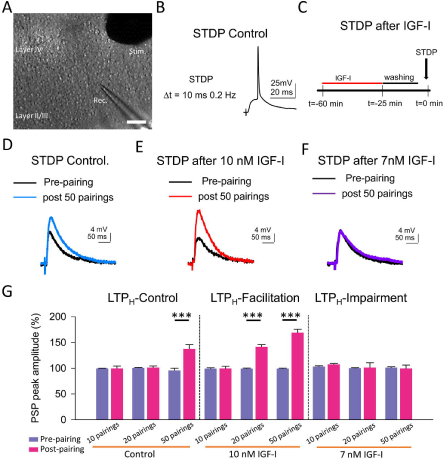
Hebbian LTP is favored and impaired by IGF-I 10nM and 7nM respectively. **A**. DIC image showing the recording (Rec.) located in layerII/III and the stimulation (Stim.) electrodes locate in Layer IV in a slice (scale bar 100 µm). **B.** Representative responses recorded (PSP followed by an AP with a 10 ms delay and a frequency of 0.2 Hz) during the STDP protcolo in control. **C.** Time course scheme showing the IGF-I exposure, washout and STDP induction (black arrow). **D.** Superimposed representative PSPs (pre-pairing, black trace) and 40 minutes after 50 pairings (post 50 pairings, blue trace) in the control experiment of STDP. **E**, same as D but when 10nM IGF-I was bath applied before the STPD. **F**, same as D but when 7nM IGF-I was bath applied before the STPD. **G.** Bar plot showing the PSP peak amplitude as percentage of the control before (pre-paring, blue bars) and 40 minutes after (post-pairing, red bars) the aplication of the STDP protocol in which 10, 20 and 50 parings were applied in control (LTP_H_-Control), or after 10nM IGF-I (LTP_H_-facilitation) or after 7nM IGF-I ( LTP_H_-impairment).

### Bidirectional modulation of synaptic transmission by IGF-I levels

We next compared the effects of these two doses of IGF-I on synaptic transmission. For these set of experiments, we recorded at L2/3 PNs (**Fig. 1A**) the postsynaptic potentials (PSPs, **Fig. 2B**) and currents (PSCs, **Fig. 2D top**) evoked by stimulation of layer 4. After 5 min of recording PSPs and PSCs, 10nM or 7nM IGF-I was applied during 35 min. As previously published ((Noriega-Prieto et al., 2021), 10nM IGF-I induced a long-term potentiation (LTP) of the PSPs (termed **LTP_IGF-1_**) that remained 30 min after IGF-I washout (from 100.70 ± 1.58 to 146.80 ± 9.06 % of amplitude at 60 min after IGF-I, P<0.01, n=8; **Fig. 2B and C, IGF-I 10nM**). However, the PSCs were transiently potentiated, returning to control values after IGF-I washout (from 98.58 ± 0.52 to 100. 29 ± 5.89 % of amplitude 60 minutes after IGF-I, P>0.05, n=9; **Fig. 2D, IGF-I 10nM, white circles**). Interestingly, 7nM IGF-I induced a long-term depression (LTD) of the PSPs (from 104.46 ± 0.82 to 70.10 ± 5.20 % of amplitude 60 minutes after IGF-I, P<0.01, n=5; **Fig. 2B and C, 7nM IGF-I**) and the EPSCs (termed **LTD_IGF-1_** from 103.78 ± 1.23 to 66. 22 ± 9.21 % of amplitude 60 minutes after IGF-I, P<0.05, n=5;, **Fig. 2D black circles and 7nM IGF-I**). Moreover, the PSCs were not modified when 5nM IGF-I was bath perfused (from 101.17 ± 0.88 to 91.95 ± 4.60 % of amplitude 60 minutes after IGF-I, P>0.05, n=6; **Fig. 2D grey circles, and IGF-I 5nM**). These results demonstrate that IGF-I can induce LTP or LTD of the PSPs at L2/3 PNs of the barrel cortex in a concentration dependent manner.

**Figure 2.**
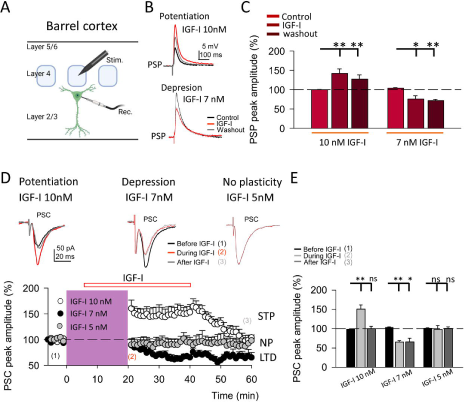
LTP and LTD of the postsynaptic potentials are induced by IGF-I 10nM and 7nM respectively. **A.** Schematic representation of layer 2/3 pyramidal neuron recording (rec) and the stimulation electrode (stim) in the barrel cortex (scheme was performed with biorender). **B top,** Superimposed representative PSPs recorded before (black trace, control), during (red trace, IGF-I) and 40 minutes after 10nM IGF-I washout (grey trace, whasout), showing the induction of PSP potentiation by IGF-I (LTP_IGF1_). **B bottom,** Superimposed representative PSPs recorded before (black trace, control), during (red trace, IGF-I) and 40 minutes after 7nM IGF-I washout (grey trace, whasout), showing the induction of PSP depresion by IGF-I (LTD_IGF1_). **C**. Bar plot showing the PSP peak amplitude as percentage of the control before (control) during (IGF-I) and 40 minutes after (washout) 10nM and 7nM IGF-I. **D top.** Superimposed representative PSCs recorded before (black trace, control), during (red trace, during IGF-I) and 40 minutes after IGF-I washout (grey trace, after IGF-I), showing the induction of PSP potentiation by 10nM IGF-I, the PSP depresion by IGF-I 7nM and the no plasticity by IGF-I 5nM. **D bottom.** Time course of the PSC peak amplitude expresed as percetenge of control before, during and after washing out the IGF-I 10nM (white circles, short term potentiation STP) 7nM (black circles, LTD) and 5nM (grey circles, no plasticity, NP). **E.** Bargraph showing the PSC change percentage showed in D bottom.

We next analyzed the locus of expression of IGF-I mediated LTD of the EPSCs. We first studied whether it was paralleled by a decrease in the probability of glutamate release. We pharmacologically isolated the EPSCs by blocking GABA_A_ inhibition with PiTX, and applied 7nM IGF-I (**Supp. Figure 1**). Under these conditions, IGF-I induced a similar LTD of the ESPCs than IGF-I 7nM, that lasted at least 40 minutes of recording (from 99.59 ± 0.44 to 67.36 ± 3.87 % of amplitude, P<0.001, n=6; **Supp. Figure 1A, black circles**). This effect was inhibited with the IGF-IR antagonist NPV-AEW 554 (from 100.37 ± 0.37 to 94.87 ± 3.67 % of amplitude, P>0.05, n=5; **Supp. Figure 1A, white circles**). To investigate the pre or postsynaptic locus of expression of this LTD, we analyzed the effect of IGF-I on the pair pulse ratio (PPR) and on the EPCS variance by measuring the coefficient of variation (CV). The effect of IGF-I on both the PPR (from 0.98 ± 0.01 to 1.28 ± 0.10 before and after IGFCI respectively, P<0.05, n=6; **Supp. Figure 1B left)** and the analysis of 1/CV^2^ plots (linear correlation R^2^=0.974, **Supp. Figure 1B right)** revealed that **LTD_IGF-1_** was due to a decrease in the probability of release of glutamate.

### Cytosolic calcium levels determine the sign of IGF-I mediated synaptic plasticity

Neurons secrete IGF-I by an activity-dependent pathway of exocytosis, and a mild depolarization is sufficient to induce IGF-I secretion in olfactory bulb neurons (Cao et al., 2011). Therefore, we next tested whether changes in cytosolic Ca^2+^ levels of postsynaptic PNs were involved in IGF-I potentiation of the EPSC. We carried out similar experiments as before, but in the presence of the Ca^2+^ chelator BAPTA (20 mM) in the recording pipette (**Fig. 3A**). Under these conditions, 10nM IGF-I induced an EPSCs depression (43.90 ± 4.04 % of baseline, P<0.001, n=6, **Fig. 3A and B**, **BAPTA**) rather than an EPSCs potentiation observed in the absence of BAPTA. This EPSC depression was abolished in the presence of NPV-AEW 554 (from 99.42 ± 1.79 to 104.08 ± 4.39 % of amplitude, P>0.05, n=5; **Fig. 3B, BAPTA+NVP**). These results indicate that IGF-I mediate EPSCs potentiation or depression depending on the cytosolic Ca^2+^ level, being all these forms of synaptic plasticity dependent on the activation of IGF-IRs.

**Figure 3.**
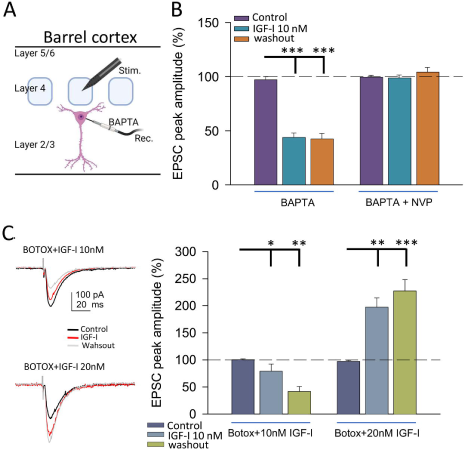
IGF-I 10nM induces LTD when postsynaptic calcium increases and endosome exocytosis are prevented by BAPTA and BOTOX respectively. **A.** Scheme showing the recorfing of layer 2/3 pyramidal neuron with a patch pipete containing 20mM Bapta and the stimulating electrode located at layer 4 of the Barrel cortex (scheme was performed with biorender). **B**. Bar plot showing the EPSC peak amplitude as percentage of the control under BAPTA in the patch pipette before (control, purple bars), during (10nM IGF-I, blue bars) and 40 minutes after washout of 10nM IGF-I (whasout, orange bars) in normal ACSF (BAPTA bars) and in ACSF containig NVP (BAPTA+NVP bars). Note that under BAPTA LTD is induced by 10nM IGF-I. **C left top.** Superimposed representative EPSCs recorded before (control, black trace), during (IGF-I, red trace) and 40 minutes after washout of 10nM IGF-I (washout, grey trace) with a BOTOX containing intracelullar solution in the patch pipette (Botox+ IGF-I 10nM). **C left bottom.** Same as top but increasing IGF-I concentration to 20mM. **C right**. Bar plot showing the EPSC peak amplitude as percentage of the control under Botox in the patch pipette before (control, darkblue bars), during (IGF-I, middleblue bars) and 40 minutes after washout of IGF-I (washout, greenbars) in normal ACSF (BAPTA bars) when applying IGF-I 10nM (left bars, Botox+10nM IGF-I) and 20nM (right bars, Botox+20nM IGF-I). Note that under Botox, LTD is induced by 10nM IGF-I and that LTP can be restored by increasing the IGF-I concentration to 20nM.

Synaptotagmin 10 (Syt10) acts as the Ca^2+^-sensor that triggers IGF-I exocytosis in olfactory bulb neurons (Cao et al., 2011). Thus, we prevented exocytosis by using the light chain of the B type botulinum toxin (i.e., Botox 0.5µM), which inhibits the SNARE protein-mediated membrane fusion of endosome complexes, and tested whether IGF-I effects depend on exocytosis. Surprisingly, 7nM IGF-I did not modulate the EPSCs (data not shown), while IGF-I 10nM depressed the EPSCs under BOTOX (58.74 ± 7.57 % of baseline, P<0.01, n=6; **Fig. 3C, BOTOX+IGF-I 10nM**), suggesting that higher IGF-I concentrations are required for LTD_IGF1_ under Botox. Interestingly, increasing IGF-I to 20nM was able to induce the potentiation of the EPSCs (130.09 ± 18.78 % of baseline, P<0.001, n=7; **Fig. 3C, BOTOX+IGF-I 20nM**) indicating that higher IGF-I concentrations are required to LTP_IGF1_ under Botox probably because it blocked the activity-dependent release of IGF-I from the postsynaptic neuron.

### Synaptic stimulation and Spiking activity can induce IGF-I mediated synaptic plasticity

Since cytosolic calcium levels and exocytosis are determinant in the induction of LTD and LTP by IGF-I, we next tested whether increases of cytosolic calcium induced by synaptic stimulation could be enough to induce IGF-I-mediated synaptic plasticity. After 5 min of recording the EPSCs, we increased the intensity of synaptic stimulation (SSI, synaptic stimulation increase) until a PSP followed by and action potential was recorded (**Fig. 4A and B**). Next, we maintained evoked these suprathreshold responses for 15 minutes by SSI, and then we turned the stimulation intensity back to control values. This protocol of stimulation induced a LTD of the EPSCs (from 101.35 ± 2.43 to 54.54 ± 8.43 % of amplitude, P<0.01, n=5; **Fig. 4A and B, black circles and bar**) that was prevented with NVP (from 101.09 ± 0.91 to 94.33 ± 4.50 % of amplitude, P>0.05, n=6; **Fig. 4A and B, white circles and bar**), or under Botox (from 100.33 ± 1.01 to 100.60 ± 2.43 % of amplitude, P>0.05, n=6; **Fig. 4A and B, red circles and bar**). These results suggest that suprathreshold responses induced by SSI are sufficient to induce an IGF-I-mediated LTD of the EPSCs dependent on endosome exocytosis. Finally, we checked whether simulation of these suprathreshold responses (**Fig. 4 C and D,** Simulated Spike) by injecting them through the patch recording electrode could induce a similar LTD of the EPSCs. As shown in figure 4 C and D, simulation of suprathreshold responses for 15 minutes was enough to induce LTD of the EPSCs (40.56 ± 3.58 % of baseline, P<0.01, n=5; **Fig. 4C and D purple circles and bar**), that is prevented with NVP (3.25 ± 3.38 % of baseline, P>0.05, n=5;

**Figure 4.**
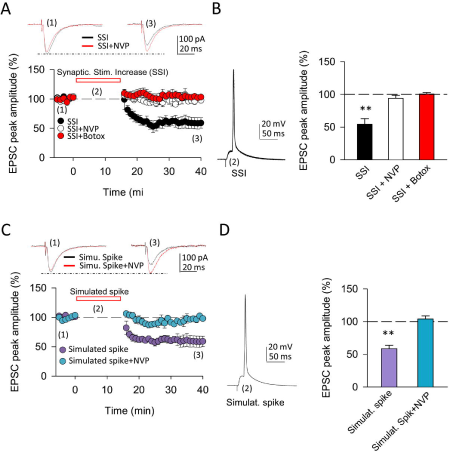
Spiking activity is able to induce LTD_IGF1_ without exogenous application of IGF-I. **A top.** Superimposed representative EPSCs recorded before (1) and 40 minutes after (3) spiking activity evoked by synaptic sitimulation increase (2, SSI) in normal ACSF (black trace, SSI) and under NVP (red trace, SSI+NVP). **A bottom**. Time course of the EPSC recorded before, during and after SSI in control (SSI, black circles), under NVP (SSI+NVP, white circles) and under Botox (SSI+botox, red circles). **B left.** Respresentative spiking activity evoked by the synaptic sitimulation increase (2). **B right.** Bar plot showing the EPSC peak amplitude as percentage of the control after SSI in ASCF ( SSI, black bar), under NVP (SSI+NVP, white bar) and under Botox (SSI + Botox, red bar). Note that SSI is able to induce an LTD that is prevented by both NVP and Botox and then is very similar to LTD_IGF1_. **C top.** Superimposed representative EPSCs recorded before (1) and 40 minutes after (3) spiking activity evoked by simulated spike (2, simulated spike) in normal ACSF (black trace, simu. spike) and under NVP (red trace, simu. spike+NVP). **C bottom**. Time course of the EPSC recorded before, during and after SSI in control (SSI, black circles), under NVP (SSI+NVP, white circles). **D left.** Respresentative simulated spike (2). **D right.** Bar plot showing the EPSC peak amplitude as percentage of the control after simulates spike in ASCF ( simulat. spike, purple bar), under NVP (simulat. spike+NVP, blue bar). Note that simulated spikes are able to induce an LTD that is prevented by NVP similar to LTD_IGF1._

**Fig. 4C and D blue circles and bar**). These results suggest that suprathreshold responses can induce increases in postsynaptic calcium levels that trigger the release of IGF-I and subsequent induction of LTD of the EPSCs.

### IGF-IR is present at presynaptic terminals of both excitatory and inhibitory synapses

We finally analyzed whether IGF-IRs were present at presynaptic terminals of excitatory in the postsynaptic L2/3 PNs. Electron micrographs demonstrated IGF-IR silver-enhanced immunogold labeling (**Fig. 5A-B**, **white arrows**) at presynaptic terminals of both excitatory (**Fig. 5A**) and inhibitory (**Fig 5B**) synapses. The presence of IGF-IR in the presynaptic terminals of both glutamatergic and GABAergic synapses supports the presynaptic LTD of the EPSCs induced by 7nM IGF-I (present results) and the presynaptic LTD of the IPSCs generated by 10nM IGF-I (Noriega-Prieto et al., 2021).

**Figure 5.**
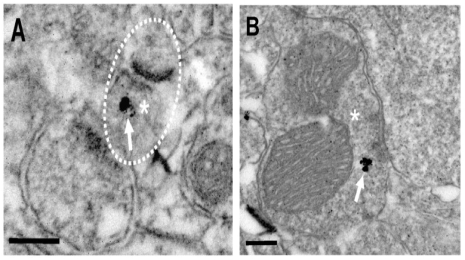
IGF-IRs are present at presynaptic sites of both excitatory and inhibitory synapses. Electron micrographs demonstrating IGF-IR silver enhanced immunogold labeling (white arrows) within both excitatory (**A**) and inhibitory (**B**) synapses. Immuno-particles were found associated to presynaptic terminals (**A and B**). Asterisks indicate presynaptic terminals. Scale bars, 0.2 μm.

## DISCUSSION

In the central nervous system, IGF-I/IGF-IR signaling is critical for experience-dependent synaptic and neuronal plasticity in sensory cortices, adult neurogenesis (LLorens-Martín et al., 2010), synaptic vesicle release, and neuronal excitability (Nuñez et al., 2003; Maglio et al., 2021). Our present results challenge the standard view that IGF-I favours cognition by inducing LTP in cortical circuits, and expands the range of brain IGF-I actions. Thus, we provide novel evidence that IGF-I levels determine the sign of plasticity induced at the barrel cortex; high levels induce LTP, while lower levels induce LTD, which in turn, favour or impair Hebbian LTP, respectively **(Fig. 6)**. Although the observation that IGF-I action on the brain is crucial for learning and memory is well stablished, here we are proposing an additional interpretation of the classical IGF-I mechanism. The results presented in this study support the importance of brain IGF-I levels as a variable in relation to synaptic plasticity, and therefore in cognitive functions.

**Figure 6.**
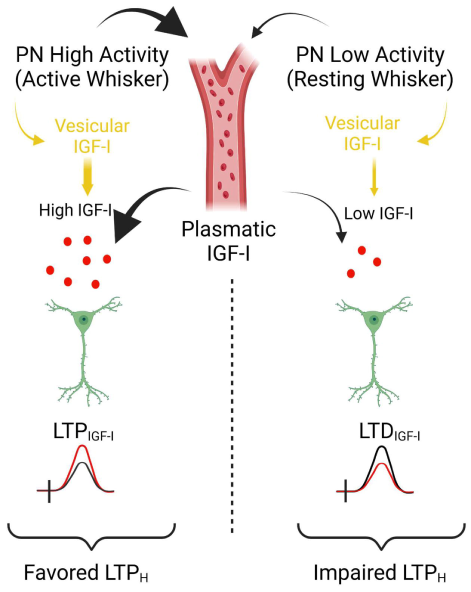
Model showing the concentration dependent actions of IGF-I. Simplified cartoon sumarizyng the novel results revealed in this work. **Left part**, under high extracellular levels of IGF-I (between 10 and 20nM) maybe obtained during high activity at pyramidal neurons (PN) of the barrel cortex during processing information of active whisker LTP_IGF1_ is induced what favours Hebbian LTP and the associatve learning and memory. Note that high extracelular IGF-I levels during spiking activity at barrel cortex are achieved by vesicular release of IGF-I stored insice PNs and maybe because facilitated IGF-I entry from plasmatic IGF-I. **Righ part**, on the contray during low activity of barell cortes PNs during resting whisker, both vesicular release of IGF-I and IGF-I entry from plasmatic levels should be lowered and then a lower IGF-Iextracelular concentation is expeted ( below 10nM) and LTD_IGF1_ is induced what impairs Hebbian LTP and the associatve learning and memory (scheme was performed with biorender).

Extracellular IGF-I activates IGF-IRs under resting conditions, maintaining basal transmission and ongoing spiking activity in the physiological range, and induces synaptic modulation (Gazit et al., 2016)). Presynaptic IGF-IRs are basally active, thus regulating glutamatergic synaptic transmission by modulating the glutamate release probability (Gazit et al., 2016). In fact, it can be concluded that tonic release of IGF-I and subsequent activation of IGF-IRs modulates synaptic vesicle release, leading to a short term depression in excitatory hippocampal neurons (Gazit et al., 2016). In agreement with these results, we also observed a long-term depression of synaptic transmission in the absence of postsynaptic activity (when firing at the recorded PN was prevented in voltage clamp) or when BAPTA abolished the cytosolic calcium increase mediated by this activity at PNs. These results strongly suggest that the tonic modulation of synaptic transmission induced by IGF-I described at the hippocampus is also present at glutamatergic synapses of the barrel cortex.

Our previous results show that 10nM IGF-I induces a long-lasting depression of the IPSCs resulting in an LTP of the PSPs, that we have termed LTP_IGFI_ (Noriega-Prieto et al., 2021). Moreover, we demonstrated that IGF-IR activation favours the induction of NMDAR-dependent LTP and improves texture discrimination of the mouse whiskers. Here we show that lower IGF-I levels (7nM) induce a long-lasting depression of the EPSCs that would result in LTD of the PSPs, that we have termed LTD_IGFI_ and impairs the induction of NMDAR-dependent LTP. This result is supported by our results showing the presence of IGF-1R in the presynaptic terminal of excitatory synapses. In addition, there is evidence suggesting this mode of action is seen in the hippocampus where IGF-I, possibly acting via GABAergic neurons, can induce the release of GABA to regulate endogenous ACh release (Seto et al., 2002). Moreover, olfactory learning in a social context selectively induces LTP of the GABAergic component of reciprocal synapses between granule and mitral cells in the medial olfactory bulb (MOB), requiring an autocrine and/or paracrine action of IGF-I to enhance postsynaptic GABA receptor function. Indeed, blocking Ca^2+^-triggered IGF-I release prevents GABAergic LTP (Liu et al., 2017). At any rate, our observations contribute to the notion that IGF-I modulates both excitatory and inhibitory synaptic activity throughout the CNS (Maglio et al., 2021; Noriega-Prieto et al., 2021). According to our results, IGF-I is able to induce a dual effect on glutamatergic synaptic transmission by the activation of IGF-IRs. In fact, both a pre-synaptically IGF-I-mediated potentiation and depression of the EPSCs were observed, and both were prevented by NPV-AEW 554 (Maglio et al., 2021; Noriega-Prieto et al., 2021)

On the other hand, we also observed that IGF-I induced an increase in glutamatergic synaptic transmission through the activation of IGF-IRs. This increase is dependent on the IGF-I-mediated increase in PNs activity, since it was absent when IGF-I was applied to voltage-clamped PNs. As discussed above, IGF-I-induced potentiation of excitatory synaptic transmission is dependent on the presence of exogenously applied IGF-I, thereby pointing to the importance of reaching specific local levels of IGF-I. Whereas there is a tonic IGF-I release that induces depression of glutamatergic synaptic transmission (Gazit et al., 2016), our results suggest that the bidirectional effect of IGF-I on the modulation of the EPSCs may depend on the levels of IGF-I reached (**Fig. 6**). An IGF-I -mediated EPSC depression is produced when PNs fires during supra-threshold responses, whereas an IGF-I-mediated EPSC potentiation is induced by bath applied IGF-I. Under physiological conditions, higher levels of IGF-I could be reached in a neuronal-activity dependent manner (Nishijima et al., 2010). In fact, we described that serum IGF-I input to the brain is regulated by an activity-driven process that includes increased blood-brain barrier permeability to serum IGF-I (Nishijima et al., 2010). Also, it is demonstrated that physical exercise induces increased brain uptake of serum IGF-I by specific groups of neurons throughout the brain (Carro et al., 2000; Nuñez et al., 2003). This increase in the uptake of IGF-I, under both mentioned physiological circumstances induces an increase in neuronal excitability, which perfectly correlates with our *in vivo* results(Noriega-Prieto et al., 2021). Our in vitro results suggest that this increase in IGF-I uptake would be essential in EPSC potentiation by increasing local IGF-I levels to those required for the modulation of synaptic transmission, playing a role in the increase in cortical activity.

In summary, our previous findings together with the present work reveal novel mechanisms of IGF-I signalling in the cortex. IGF-I induces bimodal regulation of the excitatory and inhibitory synaptic transmission depending on its levels. This bidirectional action probably contributes to favour or impair the generation of associative memories impacting on the behavioural performance of barrel cortex-related texture discrimination tasks.

## Author contributions

J.A.N.-P., L.E.M., I.T.-A., and D.F.d.S. designed research; J.A.N.-P., L.E.M., J.C.D., and A.G. performed research; J.A.N.-P., L.E.M., J.C.D., and A.G. analysed data; J.A.N.-P., L.E.M., and D.F.d.S. wrote and reviewed the manuscript. J.A.N.-P., L.E.M., J.C.D., A.G., I.T.-A., and D.F.d.S. reviewed drafting of the manuscript, the final version of which they approved for submission.

## ACKNOWLEDGMENTS

This work was supported by the following Grants: PID2020-119358GB-I00 to D. Fernández de Sevilla and PID2019-104376RB-I00 (AEI/FEDER, UE) to I. Torres-Alemán. We would like to thank Ms. Marta Callejo for excellent technical assistance.

---

**Supplementary Figure 1.**
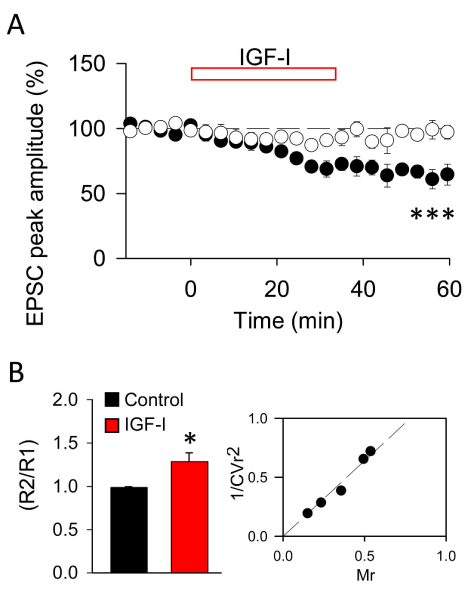
LTD_IGF1_ is mediated by a long-term decrease in the glutamate release probability. A. Time course of the EPSC recorded under PiTX before, during and after washing out IGF-I 7nM in control (black circles, LTD_IGF1_) and under NVP (white circles). **B left**. summary data showing the paired-pulse ratio (R2/R1) of EPSCs recorded before (black bar, control) and during IGF1 7nM (red bar, IGF-I). Note the increase in the paired-pulse ratio suggeting an incresae in the glutamate probability of release. **B Right**.Plot of the variance (1/CV2, where CV is coefficient of variation) as a function of the mean peak EPSC amplitude in the presence of IGF-I normalized to control conditions (Mr). Note that the values were grouped following the diagonal suggesting that LTD_IGF1_ was due to a change in presynaptic release properties.

